# pan-ECM: live brain extracellular matrix imaging with protein-reactive dye

**DOI:** 10.1101/2023.03.29.534827

**Authors:** Xiaoqian Ge, Xueqi Xu, Qi Cai, Hejian Xiong, Xie Chen, Yi Hong, Xiaofei Gao, Yao Yao, Robert Bachoo, Zhenpeng Qin

**Affiliations:** Department of Mechanical Engineering, The University of Texas at Dallas, Richardson, Texas, United States; Department of Bioengineering, The University of Texas at Arlington, Arlington, Texas, United States; Department of Internal Medicine, University of Texas Southwestern Medical Center, Dallas, Texas, United States; Department of Molecular Pharmacology and Physiology, Morsani College of Medicine, University of South Florida, Tampa, Florida, United States; Department of Neurology and Neurotherapeutics, University of Texas Southwestern Medical Center, Dallas, Texas, United States; Harold C. Simmons Comprehensive Cancer Center, University of Texas Southwestern Medical Center, Dallas, Texas, United States; Department of Bioengineering, Center for Advanced Pain Studies, The University of Texas at Dallas, Richardson, Texas, United States; Department of Surgery, The University of Texas at Southwestern Medical Center, Dallas, Texas, United States; Center for Advanced Pain Studies, the University of Texas at Dallas, Richardson, Texas, United States

## Abstract

The brain extracellular matrix (ECM), consisting of proteins and glycosaminoglycans, is a critical scaffold in the development, homeostasis, and disorders of the central nervous system (CNS) and undergoes remodeling in response to environmental cues. Live imaging of brain ECM structure represents a native view of the brain ECM but, until now, remains challenging due to the lack of a robust fluorescent labeling approach. Here, we developed a pan-ECM method for labeling the entire (Greek: pan) brain ECM network by screening and delivering a protein-reactive dye into the brain. pan-ECM enables imaging of ECM compartments in live brain tissue, including the interstitial matrix, basement membrane (BM), and perineuronal nets (PNNs), and even the ECM in glioblastoma and stroke mouse brains. This approach provides access to the structure and dynamics of the ECM and enhances our understanding of the complexities of the brain ECM and its contribution to brain health and disease.

## Introduction

Neurons and glia in the brain secrete molecules to form a three-dimensional ECM scaffold that binds to specific ECM receptors on cell surfaces,^1-3^ with alterations in ECM scaffolds implicated in numerous neurological disorders including Alzheimer’s disease, Parkinson’s disease, and epilepsy.^2, 4-7^ Visualizing the brain ECM allows quantifying changes in the ECM organization in response to various stimuli, such as injury, disease, and developmental cues. For decades, the standard method of visualizing brain ECM involves immunostaining individual ECM components including hyaluronic acid (HA) and chondroitin sulfate proteoglycans (CSPG) in fixed tissues.^7, 8^ However, chemical fixation and dehydration processes in immunostaining lead to substantial structural shrinkage and do not provide a native view of the brain ECM.^9^ Cryo-fixation electron microscopy better preserves the extracellular structure and has provided important insights into the brain extracellular space (ECS) with high spatial resolution, but it does not allow for real-time imaging of brain tissue due to the necessity of prior fixation.^10^ Second harmonic generation (SHG) allows live imaging of fibrillar collagen,^11-14^ one of ECM components, but collagen is only distributed in the vasculature.^15, 16^ Recent advancements in single-particle tracking and super-resolution shadow imaging (SUSHI) of freely diffusing dyes in live brain tissue have become widely used for studying complex biological activities in the ECS.^17-21^ However, there are currently only a limited number of tools and methods available for visualizing, identifying, and mapping ECM networks in live brain tissue.

We have developed a method called pan-ECM that enables live visualization of the ECM network in various brain regions. Specifically, we screened an N-hydroxysuccinimide ester (NHS)-tagged dye that reacts with the ECM proteins through the amine-reactive crosslinker chemistry with minimal dye aggregation or cellular uptake. After stereotaxic delivery, the dye conjugates uniformly on the interstitial matrix, perineuronal nets (PNNs), and basement membrane (BM), which are three major compartments of the brain ECM.^1^ The pan-ECM approach allows for imaging the brain ECM structures and analyzing the volume fraction of the brain ECM across different brain regions (such as the cortex, hippocampus, striatum, and cerebellum). Moreover, we demonstrated that pan-ECM can visualize the ECM structures in mouse stroke and glioblastoma tissue and reveals that a significant disruption of the ECM in stroke tissue and a dense and complex ECM in glioblastoma. The pan-ECM method represents a significant breakthrough, enabling visualization of the ECM in live brain tissue that was previously difficult, and is expected to enhance our comprehension of the ECM’s role in physiological and pathological conditions.

## Results

### Dye screening for pan-ECM labeling

First, we performed a two-step dye screening for pan-ECM labeling (**Fig. 1a**). The screening involves firstly screening dyes without significant dye aggregation in the ECS (Step 1) and then in vivo testing for minimal endocytosis and maximal diffusion to label a large ECM volume in the brain (Step 2). For Step 1, we selected eight widely used dyes that have carboxylic acid group and prepared acute hippocampal brain slices to incubate with dye-filled artificial cerebrospinal fluid (ACSF) solution (**Supplementary Table 1**). As the hippocampus has large neuronal soma and dense axons and dendrites, we imaged this area to check for any dye aggregation using a spinning disk confocal microscope. The imaging depth for acute hippocampal brain slices is at least 30-50 µm to avoid the detection of dead cells that are highly dye permeable. Confocal images showed that four hydrophilic dyes, Atto 488, calcein, Cy3b, and Atto 655 (**Fig. 1b**), exhibited uniform distribution in the hippocampal ECS. However, two hydrophilic dyes (Atto 565 and Alexa 647) displayed moderate aggregation and two hydrophobic fluorophores (Cy5 and Atto 647N) exhibited a high degree of aggregation, as fluorophores interact lightly or heavily with the brain ECM via the hydrophobic-hydrophobic interaction. Considering the low photostability of calcein,^22^ we selected non-aggregated dyes, Atto 488, Cy3b, and Atto 655, for further evaluation as pan-ECM candidates due to their high quantum yield, brightness, and photostability.^23^

**Fig. 1.**
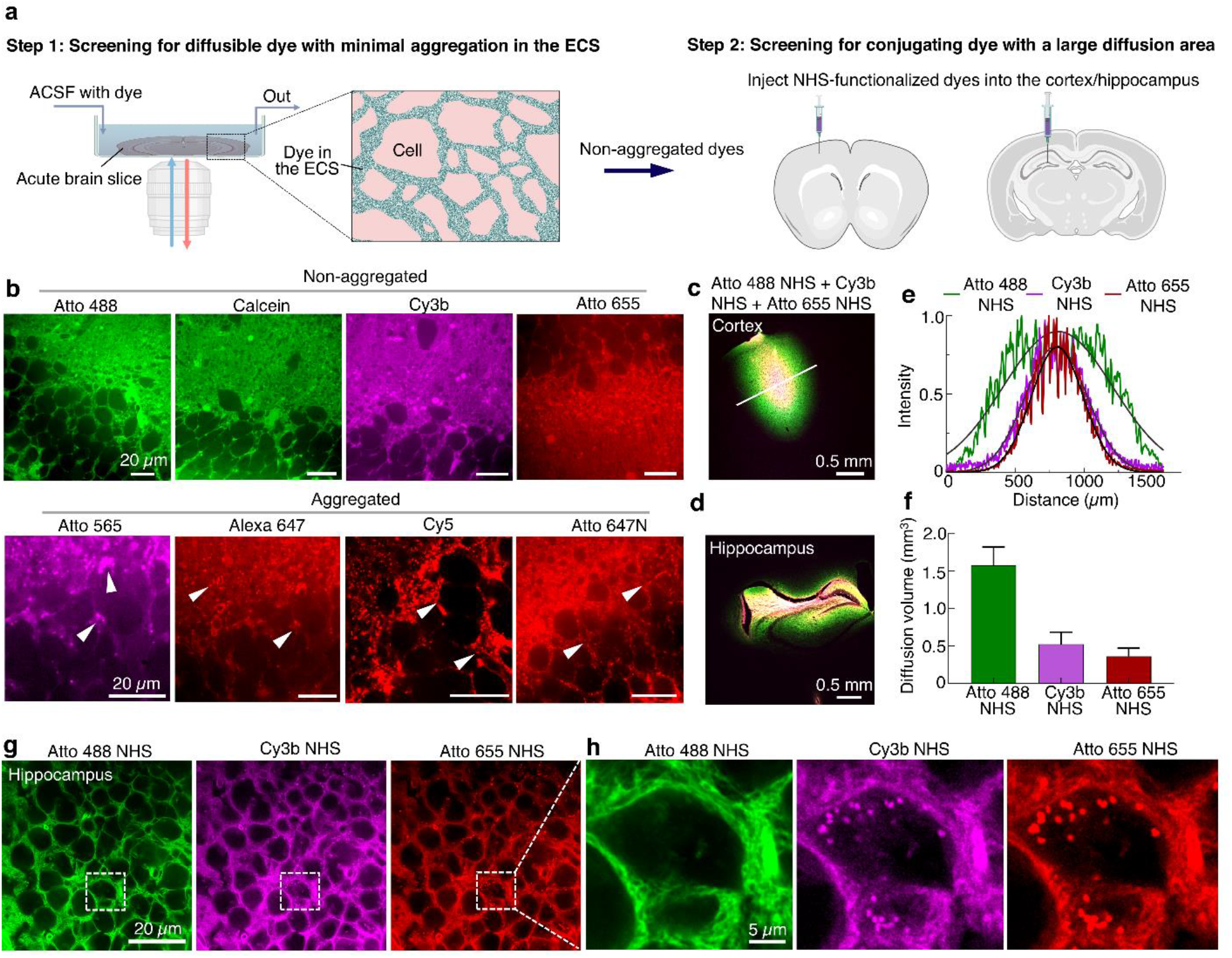
Dye screening for pan-ECM labeling with minimal aggregation and large diffusion area. (**a**) Workflow for the dye screening for the pan-ECM labeling. (**b**) Representative confocal images for screening diffusible dyes with minimal aggregation in the hippocampal ECS. Dye aggregation was marked with white arrows. (**c-d**) Large field-of-view images of the extracellular structures in the brain cortex (c) and hippocampus (d), co-labeled with Atto 488 NHS, Cy3b NHS, and Atto 655 NHS. Each dye concentration was controlled at 0.17 mM, and imaging was performed on fixed brain slices with 30 µm thickness. (**e**) Fluorescence intensity profiles across the white line in panel c. Black lines are Gaussian fitting of intensity profiles. (**f**) The in vivo diffusion volume of Atto 488 NHS, Cy3b NHS, and Atto 655 NHS (N = 3 mice, mean ± SD). Star indicates P value of a one-way ANOVA test; ***: P < 0.001. (**g**) High-resolution confocal images of the extracellular structure in the hippocampus, co-labeled with Atto 488 NHS, Cy3b NHS, and Atto 655 NHS. (**h**) Closed-up images of dashed boxes in panel g.

Further screening of the pan-ECM candidate was conducted by examining the in vivo labeling and the diffusion properties (**Fig. 1a** Step 2). We tested three dyes with the NHS group, which conjugates with proteins in the brain ECM network upon injection while diffusing around the injection site. Dyes that spread over longer distances would result in a lower dye concentration at the injection site, reducing the likelihood of cellular uptake or endocytosis. The mixture of Atto 488 NHS, Cy3b NHS, and Atto 655 NHS was stereotaxically injected into the cortex and hippocampus with the same concentration. The comparison shows that Atto 488 NHS diffused longer distance and has 3- to 4-fold higher diffusion volume than Cy3b NHS and Atto 655 NHS (**Fig. 1c-f**, and **Supplementary Fig. 1**). Further, high-resolution confocal images show that Cy3b NHS and Atto 655 NHS dyes appear in the hippocampus neurons via endocytosis, but not with Atto 488 NHS (**Fig. 1g-h**). Based on these results, Atto 488 NHS was selected for pan-ECM labeling and imaging.

### pan-ECM labels three major compartments of the brain ECM without cellular uptake

Next, we examined whether pan-ECM labels three major compartments of the brain ECM, namely the interstitial matrix, BM, and PNNs (**Fig. 2a**). To probe the interstitial matrix, we compared pan-ECM labeling with immunostaining of HA and CSPG. Confocal images showed that hyaluronan binding protein (HABP) staining homogeneously colocalizes with pan-ECM labeling (**Fig. 2b** and **Supplementary Fig. 2a, c, e**, and **g**). Furthermore, Pearson’s correlation coefficient (PCC) was used to quantify the colocalization between pan-ECM labeling and HABP staining, exhibiting a reasonable correlation (>0.5) in four different brain regions including the cortex, hippocampus, caudate-putamen and cerebellum (**Supplementary Fig. 2b, d, f**, and **h**). Similar to HABP staining, the CSPG staining also exhibited significant overlap with the pan-ECM labeling (**Fig.2c**).

**Fig. 2.**
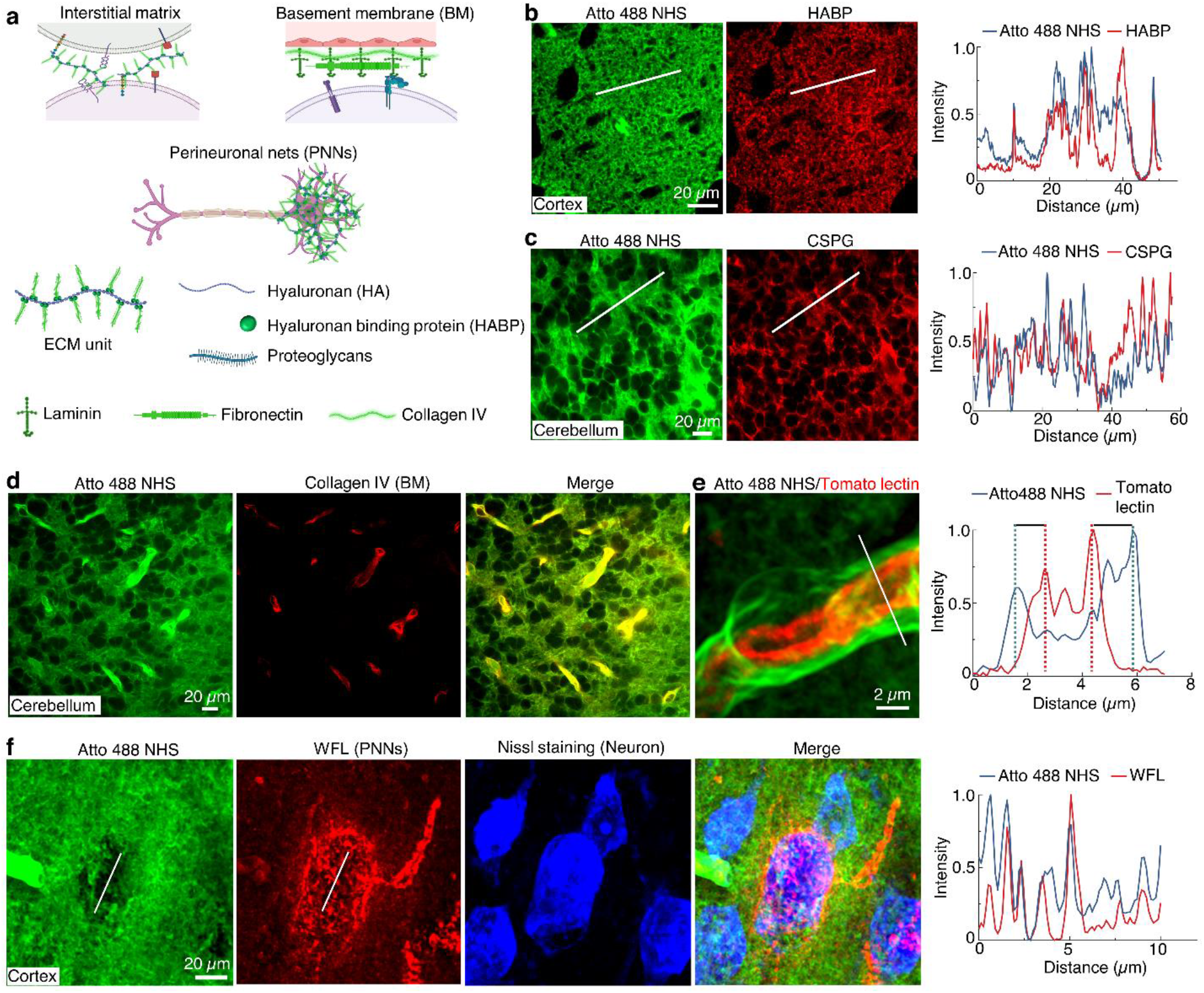
pan-ECM broadly labels the ECM, including the interstitial matrix, basement membrane (BM), and perineuronal nets (PNNs). (**a**) Schematic showing of three major distribution of ECM, interstitial matrix, BM, and PNNs; single ECM unit and its components, including HA, HABP, proteoglycans, laminin, fibronectin, and collagen IV. (**b-c**) Left: Confocal images of Atto 488 NHS labeled structures with HABP (b) and CSPG staining (c); Right: fluorescence intensity profiles along the white lines in panel b and c. (**d**) Confocal images of Atto 488 NHS labeled structures, and collagen IV staining. (**e**) Left: merged image of the Atto 488 NHS labeled structures in the cortex and the vascular lumen labeled with tomato lectin-Dylight 649; Right: intensity profiles along the white lines in panel f, suggesting Atto 488 NHS labeling presents outside of the vascular lumen. (**f**) Confocal images of Atto 488 NHS labeled structures, WFL staining, and Nissl staining; both images were obtained from the maximum-intensity projection of a Z-stack.

We further characterized pan-ECM labeling of BM and PNNs. We stained the collagen IV protein, one of the components in the BM. Across four brain regions (cortex, cerebellum, hippocampus, and caudate-putamen), collagen IV staining located on the brain vasculature and overlapped well with pan-ECM labeling (**Fig. 2d** and **Supplementary Fig. 3**). We reason that by intraparenchymal injection, pan-ECM labels the abluminal side of the vasculature, rather than the luminal surface. To test this, we intravenously (i.v.) injected the mice with tomato lectin-Dylight 649, which labels the luminal surface, in addition to the pan-ECM labeling. A comparison of the two labeling shows that Atto 488 NHS labeling has peaks outside the vascular lumen (tomato lectin-Dylight 649) (**Fig. 2e** and **Supplementary Fig. 4a**). As a control, we injected Atto 488 dye (without protein conjugation) into the hippocampus and monitored dye deposition in blood vessels 1 hour and 24 hours after injection. Our results showed no detectable dye deposition in blood vessels, indicating minimal nonspecific binding or clearance via the glymphatic system (**Supplementary Fig. 4b**). To verify if pan-ECM labels PNNs, Atto 488 NHS-labeled brain slices were stained with wisteria floribunda lectin (WFL) and NeuroTrace (Nissl staining). WFL specifically targets the sugar chains of CSPGs present in the ECM surrounding neurons, while Nissl staining labels the Nissl substance within neuronal cells. Our results showed that the WFL staining exhibited a net structure outside of the Nissl staining, thus demonstrating the localization of PNNs (**Fig.2f**). Furthermore, we observed a uniform colocalization between pan-ECM labeling and WFL staining in the cortex through line intensity analysis between these two channels (**Fig.2f**). These findings provide evidence that pan-ECM labels PNN in brain tissue.

We also assessed whether brain cells, including neurons, astrocytes, and microglia, take up Atto 488 NHS. To investigate the uptake of Atto 488 NHS in neurons, we used CaMKII-Cre;Ai14 mice, which express red fluorescent protein under the calcium/calmodulin-dependent protein kinase II (CaMKII) promoter in excitatory neurons.^24^ We stereotaxically injected Atto 488 NHS into the hippocampus of these mice, and acute brain slices were prepared for two-color live imaging after 0.5 h of injection. Our results show that Atto 488 NHS labeled structure has minimal overlap with neurons (**Supplementary Fig. 5**). To examine the cellular uptake of Atto 488 NHS in astrocytes, we injected the dye into the corpus callosum, which contains abundant protoplasmic astrocytes.^25^ Immunostaining of glial fibrillary acidic protein (GFAP) of astrocytes in Atto 488 NHS labeled fixed slice shows that astrocytes did not uptake the dye (**Supplementary Fig. 6a-c**). Similarly, staining brain slice with ionized calcium-binding adapter molecule1 (Iba1) antibody revealed no uptake of Atto 488 NHS by microglia (**Supplementary Fig. 6d**). These findings suggest that Atto 488 NHS has minimal cellular uptake in neurons, astrocytes, and microglia. Furthermore, we found cellular uptake at 24 hours and 7 days after injection, while there is no uptake in the acute phase (1-hour, **Supplementary Fig. 7**).

### pan-ECM provides wash-resistant labeling compared with diffusible dyes

We compared pan-ECM with freely diffusible dyes that are currently used for ECS labeling. To this end, Atto 488 and Atto 488 NHS dyes were injected into each side of the hippocampus (**Fig. 3a**) and subsequently analyzed confocal images. Our findings demonstrate that the labeling of the ECM in the hippocampus with Atto 488 was weak due to dye diffusion, whereas Atto 488 NHS labeling provided clear ECM structures (**Fig. 3b**). Furthermore, the intensity of the confocal images obtained with Atto 488 labeling was found to be 20-fold weaker than those obtained with Atto 488 NHS labeling (**Fig. 3c**). We also investigated the washability of Atto 488 NHS labeling. Following Atto 488 NHS labeling of the corpus callosum, we prepared acute brain slices for imaging. An extended 20 min ACSF buffer perfusion shows stable Atto 488 NHS conjugation and demonstrates the wash-resistant feature (**Supplementary Fig. 8**).

**Fig. 3.**
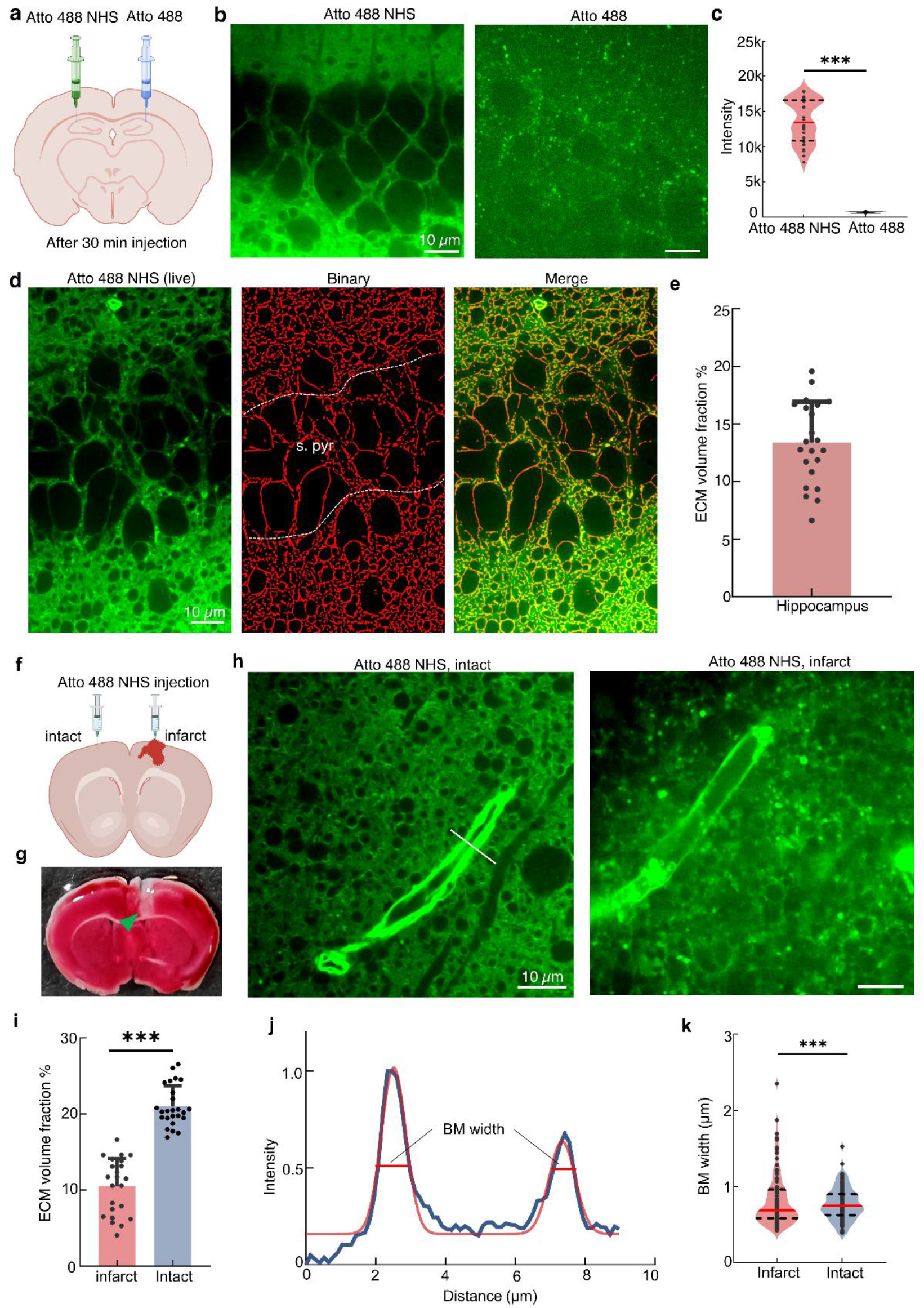
Live imaging and quantitative analysis of ECM architecture and volume fraction in healthy and stroke brain tissues. (**a**) Schematic showing of dyes injection into both hippocampus. (**b**) Confocal images of extracellular structures with Atto 488 NHS and Atto 488 labeling after 30 min injection of Atto 488 NHS and Atto 488. (**c**) Violin plot of fluorescence intensity of confocal images obtained with Atto 488 NHS, showing 20-fold fluorescence enhancement over Atto 488 labeling. Each dot represents the mean fluorescence intensity of one image. Red line, median value; dashed black lines, first quartiles (25th percentile) and third quartiles (75th percentile); n = 30 images from 6 brain slices, 3 mice; star indicates P value of a Wilcoxon signed rank test; ***: P < 0.001. (**d**) (Left) confocal image of the brain ECM of the hippocampus, labeled with Atto 488 NHS; (middle) the binary of the confocal image, indicating the stratum pyramidal (s. pyr) has a smaller volume fraction across the hippocampus; (right) the merge of the confocal and binary images. (**e**) The ECM volume fraction distribution of the hippocampus. N = 22 images from 3 mice, mean ± SD. (**f**) Schematic illustration of stereotactic injection of Atto 488 NHS to the intact and infarct area of stroke mouse. (**g**) Digital photograph of 2 mm thick brain slice after 15 min soaking in 2,3,5-triphenyltetrazoliumchloride (TTC) saline solution. The stroke area became white in the cortex, as indicated by a green arrow. (**h**) Live confocal imaging of ECM in intact and infarct areas. (**i**) The ECM volume fraction of the infarct and intact area. (**j**) Fluorescence intensity profiles are plotted from the white line in panel h. The black line is the Gaussian fitting of the intensity profiles, and the full width at half maximum (FWHM) value is considered the BM thickness. (**k**) Violin plot of the BM thickness distribution of the intact and infarct area. Red line, median value; black dashed lines, first quartiles (25th percentile) and third quartiles (75th percentile); n = 50 images from 6 brain slices; star indicates P value of a Wilcoxon signed rank test; ***: P < 0.001.

### pan-ECM enables ECM volume fraction analysis and visualization of BM changes in stroke mice

The ECM volume fraction is an essential parameter that impacts substance diffusion in the nervous tissue. To determine the ECM volume fraction (α), an image analysis approach was employed, which involved calculating the ratio of the number of pixels in the binarized structure (red color in **Fig. 3d**) to the total number of pixels in the image. The α value of the hippocampus exhibited regional variation, with an overall value of 13 ± 4% (**Fig. 3e**). Furthermore, the α was calculated as 15 ± 3% for the caudate-putamen, 16 ± 2% for the cerebellum, and 21 ± 3% for the cortex (**Supplementary Fig. 9**), and these values are comparable with previous reports for ECS volume fraction.^18, 26^

We employed the pan-ECM approach to investigate changes in the ECM following a stroke. We utilized the rose bengal stroke model and subsequently injected Atto 488 NHS into both the infarct and intact (control) areas after 24 hours of stroke (**Fig. 3f**). The stroke model was confirmed via 2,3,5-triphenyltetrazoliumchloride (TTC) staining, whereby the infarct area in the cortex appeared white (**Fig. 3g**). By comparing the ECM changes in the infarct area with the intact area, we observed a significant deterioration in the ECM structure in the infarct area, transitioning from a continuous to a discrete structure as compared to the intact area (**Fig. 3h**). To quantify these changes, we measured the ECM volume fraction after stroke and found a reduction from 22 ± 3% to 10 ± 4% for the intact area (**Fig. 3i**). Furthermore, we examined the impact of stroke on the BM changes, which can occur due to proteolytic degradation triggered by various oxidative and inflammatory responses. By plotting the fluorescence intensity profile across the BM of blood vessels with visible lumens, we determined the BM width using the FWHM obtained from the Gaussian fitting of the intensity profile (**Fig. 3j**). Our data revealed a statistically significant decrease in BM width in the infarct area (median 0.69 µm, IQR 0.58-0.96 µm) as compared to the intact area (median 0.75 µm, IQR 0.62-0.90 µm) (**Fig. 3k**). In summary, our findings using the pan-ECM approach provide valuable insights into the ECM and BM alterations following a stroke.

### pan-ECM enables visualization of glioma ECM

Lastly, we employed pan-ECM to examine glioma ECM, which undergoes significant transformation compared with healthy brain regions. To do this, we first prepared an orthotopic glioma model by implanting tdTomato fluorescence-tagged 73c glioma cells into mice brains (**Fig. 4a**). After three weeks, we intratumorally injected Atto 488 NHS to label glioma extracellular structure, as evidenced by large field-of-view images (**Fig. 4b**). To confirm that the glioma ECM was uniformly labeled with the Atto 488 NHS, we immuno-stained four ECM-related components, HABP, collagen IV, tenascin-C, and fibronectin. **Fig. 4c** shows that the glioma displays similar levels of HABP expression but higher levels of collagen IV, tenascin-C, and fibronectin than healthy brain regions. High-resolution confocal images show that both HABP and collagen IV staining colocalizes well with the Atto 488 NHS labeled structures (**Fig. 4d-g**). The tenascin-C and fibronectin distributions in the glioma were is less dense, however, they exhibit good co-localization with pan-ECM labeling (**Supplementary Fig. 10**). Staining with GFAP and Iba1 antibodies suggest no significant uptake of Atto 488 NHS by astrocytes and microglia cells (**Supplementary Fig. 11**), which coexist with glioma cells and plays a prominent role in the progression of glioblastoma.^27^ Significantly, live glioma ECM imaging indicate that the ECM in the glioma mass exhibits a lower level of complexity than the ECM present in the healthy brain regions (**Fig. 4h** and **i**). This disparity could potentially be attributed to the alteration of neuronal structures, which may be lost as a consequence of the glioma aggressive growth.^28^ These observations are crucial in unraveling the underlying mechanisms that contribute to the formation and progression of gliomas, and may help identify potential targets for therapeutic interventions.^29^

**Fig. 4.**
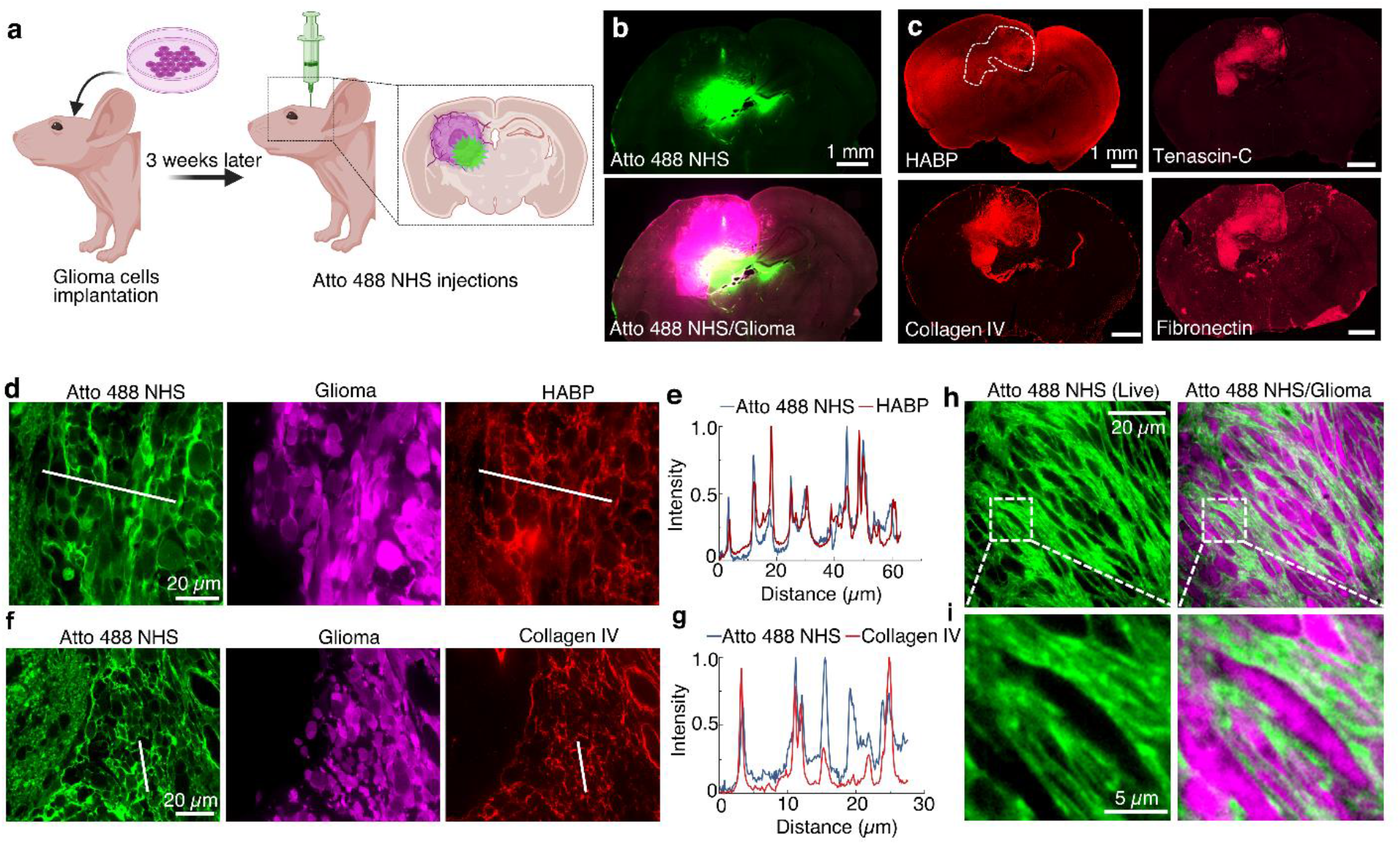
pan-ECM enables visualization of glioma ECM. (**a**) Schematic illustration of glioma model and pan-ECM labeling. (**b**) Large field-of-view images reveal the glioma and the pan-ECM labeled area. (**c**) Large field-of-view images of immunostaining of major glioma ECM components, including HABP, collagen IV, tenascin-C, and fibronectin, showing different expression levels. (**d** and **f**) Confocal images of the Atto 488 NHS labeling, glioma cell, and HABP or collagen IV staining; (**e** and **g**) Fluorescence intensity profiles along the white lines in green and red channels, showing good colocalization between Atto 488 NHS and HABP or collagen IV channels. (**h**) Live confocal imaging of the ECM in the glioma mass, Atto 488 NHS labeling, glioma cell. (**i**) The closed-up of dashed boxes in the panel h.

## Discussion

Through their cell surface receptors, neuronal and glial cells enmesh themselves in the ECM meshwork. The ECM architecture can be remodeled or altered as the brain develops, matures, or undergoes pathological changes. However, elucidating the precise topology and dynamics of ECM in response to brain activity remains a daunting task, due to the absence of a reliable labeling method. Here, we present a novel approach, pan-ECM, that enables real-time visualization and analysis of the ECM meshwork in live brain tissue, which we have validated in both healthy and diseased brains. Our method can provide an unprecedented insight into the dynamic and complex ECM network, paving the way for a better understanding of brain function in health and disease.

pan-ECM is easily implanted using stereotactic injections and can be completed within several hours, including dye injection, slicing, and imaging. Combined with brain slice culture, the pan-ECM allows visualizing the live acute brain slice and tracking dye-labeled ECM within the slice over hours. Compared with the SHG method, the pan-ECM method labels the entire network of ECM, while the SHG allows only the imaging of fibrillar collagen, which is scarce in the ECM of healthy mice brains but abundant in the ECM of gliomas (**Fig. 2d** and **Fig. 4c**). The SHG method must be combined with two-photon fluorescence; however, the pan-ECM is adaptable to standard fluorescence microscope by using a fluorescent dye. Therefore, the pan-ECM would greatly expand the applications related to the brain ECM imaging, and thus provide detailed information on the ECM topology.

pan-ECM labels the non-diffusible matrix proteins while the ECS labeling utilizes diffusible dyes. Prior work has validated the global ECS volume fraction over a large area, ∼20%, using the real-time tetramethylammonium (RTI-TMA) method,^30, 31^ and recent reports using diffusible dyes reports similar and brain region-dependent values. Using the pan-ECM, we calculated the ECM volume fraction to be around 15-20% (**Fig. 3d-e** and **Supplementary Fig. 9**), which is comparable to the ECS volume and varies across brain regions. Future work may explore the diffusible versus non-diffusible regions using a combination of ECM and ECS imaging.

One limitation of pan-ECM labeling is that it labels all proteins exposed to the extracellular environment. Thus it may attach to various cell membrane receptors, including both ECM-related and non-ECM-related receptors. The ECM-related receptors, such as integrins, syndecans, CD44, and dystroglycan,^32^ play a crucial role in cell adhesion, migration, proliferation, and differentiation. However, current imaging techniques have limited resolution and cannot effectively distinguish the ECM and membrane proteins. This issue may be resolved by developing advanced nanoscopy and appropriate receptor labeling methods.

Going forward, we believe the pan-ECM method can be used to image ECM in a broader range of disease models, including Alzheimer’s and Parkinson’s diseases. The pan-ECM technology is useful in examining the brain ECM remodeling associated with these diseases. In our study, the imaging resolution was insufficient to reveal the ultrastructural characteristics of ECM. We anticipate this pan-ECM approach, together with super-resolution microscopes, such as stimulated emission depletion (STED) or 2-photon STED,^33, 34^ could serve as an effective method for unraveling detailed ECM structures and exploring brain ECM-related scientific issues.

### Online Methods

#### Animals

Animal protocols were followed according to guidelines established by the National Institutes of Health (NIH) and approved by the Institutional Animal Care and Use Committee of The University of Texas at Dallas or University of Texas Southwestern Medical Center. Six-week-old male C57BL/6 mice were purchased from the Charles River Laboratory. Six-week-old female NU/J mice (Foxn1^nu^) were purchased from the Jackson Laboratory. Transgenic mice CamKǁ-Cre;Ai14 were bred in Dr. Robert Bachoo’s laboratory at the University of Texas Southwestern Medical Center.

Before brain surgeries, mice were deeply anesthetized using 2.5 % isoflurane vapor and placed in a stereotaxic frame via ear bars. A heating pad connected to a rectal probe kept the mice’s temperature around 34 °C. Before surgery, mice were subcutaneously administered 1 mg/kg of buprenorphine (ZooPharm) to relieve pain and administered dexamethasone (Sigma-Aldrich) at the dosage of 5 mg/kg to prevent brain inflammatory response.

#### Glioma cell line and mice injection

Mouse astrocyte-derived glioma cell line 73c (tdTomato tagged) was provided by Dr. Robert Bachoo’s laboratory.^35^ 73c cells were cultured in Dulbecco’s Modified Eagle Medium (DMEM, Thermo Fisher Scientific, catalog NO. 10569010), containing 10% fetal bovine serum (FBS, Thermo Fisher Scientific, catalog NO. A3160502), to reach ∼80% confluence before transplantation. Cells were trypsinized and resuspended in Hank’s Balanced Salt Solution (HBSS) with a density of 2 × 10^5^/μL. 0.3 μL cells/HBSS suspension were transplanted to the left hemisphere of NU/J mice (Foxn1^nu^) via a 3 mm diameter cranial window, centered at anterior/posterior (A/P) -1 mm, medial/lateral (M/L) +0.5 mm, and dorsal/ventral (D/V) 2 mm. The exposed skull was covered with Body Double™ Standard Set (Smooth-On, Inc) for protection. Next, mice were housed in the animal room for three weeks to allow glioma growth.

#### Stroke mice preparation

The photothrombotic stroke was induced as previously described with some modifications.^36^ Briefly, a ∼5 mm × 5 mm area on the exposed skull was marked with an ultrafine mark pen, centrated at A/P +1.0 mm, M/L +1.5 mm to bregma. The skull in this marked area was thinned using a dental drill, followed by positioning a fiber-coupled LED (595 nm, M595F2, *Thorlabs*) in the center. 5 min before illumination, 100 µL fresh Rose Bengal (*Sigma-Aldrich*, product No. 330000) saline solution (20 mg/mL) was intravenous (i.v.) administered to mice, and the thinned area was subsequently illuminated for 15 min. The output power of the 595 nm illumination light was set as 1.1 mW by measuring the fiber tip with a PowerMeter. Then, the exposed skull was covered with Body Double™ Standard set (Smooth-On, Inc), and mice were housed for 24 h before following experiments.

The 2,3,5-triphenyltetrazolium chloride (TTC, Sigma-Aldrich, product No. T8877) staining was adopted to visualize the stroke area.^37^ In detail, the brain of the stroke mice was extracted from and submerged in 1x PBS in a brain matrices slicer (World Precision Instrument, RBMS-600C). A 2 mm coronal brain slice was cut using two blades, covering the stroke area. Then, the brain slice was immersed in 2% TTC saline solution at 37 °C for 15 min. The stroke area was white, while other cortex regions exhibited red color.

#### Brain extracellular matrix (ECM) fluorescently labeling

Stereotaxic injections of Atto 488 NHS and other dyes were conducted to label the brain ECM of different mice. The Atto 488 NHS dye was prepared as 5 mM in dimethyl sulfoxide (DMSO), accompanied by assigning 0.5 µL dye/DMSO solution in one aliquot for -20 °C storage. Before each injection, 9.5 µL 1x phosphate-buffered saline (PBS) was added to one aliquot to prepare the new Atto 488 NHS dye solution (0.25 mM). The dye was injected using the Nanoliter 2020 Injector (World Precision Instrument) with a 30 µm diameter pipette generated by the Micropipette Puller (Sutter Instrument Co., P-97). The injection rate was controlled at 1 nL/s.

##### Fluorescently labeling of brain ECM of wild-type C57BL/6 mice

The wild-type C57BL/6 mice’s ECM positions were labeled, including the cerebral cortex, caudate-putamen, hippocampus, and cerebellum. There was only one cranial window opened in one mouse, centered at A/P +1.0 mm, M/L +1.5 mm for cerebral cortex; or A/P +1.0 mm, M/L +2.0 mm for caudate-putamen; or A/P -1.8 mm, M/L +1.3 mm for hippocampus; or A/P -6.5 mm, M/L 0 mm for the cerebellum, respectively. All coordinates were calculated from the bregma point. Roughly, 0.4 µL of fresh Atto 488 NHS dye in 1x PBS (0.25 mM) was administrated to the cerebral cortex (D/V 1 mm), 0.8 µL was injected into the caudate-putamen (D/V 3.2 mm), hippocampus (D/V 1.8 mm), or cerebellum (D/V 2.0 mm). The mouse was left in the stereotaxic frame for 0.5 hours under anesthesia with 2.5 % isoflurane treatment. Then, the mouse was ready for the acute brain slice preparation.

Atto 488 NHS, combined with Cy3b NHS and Atto 655 NHS, was used to inject the cortex or hippocampus. Cy3b NHS and Atto 655 NHS were also prepared as 5 mM in DMSO, similar to Atto 488 NHS storage and preparation described above. Before injection, Atto 488 NHS, Cy3b, and Atto 655 NHS were mixed, and each dye’s final concentration was controlled at 0.17 mM. A dye mixture of 0.4 µL was injected into the cortex, and a dye mixture of 0.8 µL was injected into the hippocampus, according to the exact coordinates.

##### Fluorescently labeling brain ECM of neurons transgenic mice

Transgenic mice were used for two-color imaging between ECM and neurons. In the CamKII-Cre:Ai14 mouse, ECM in the hippocampus was labeled using a similar method to labeling the ECM of the hippocampus of wild-type C57BL/6 mice

##### Fluorescently labeling ECM of glioma

Multiple intratumorally injections were used to label the ECM of the glioma. The previous cranial window for glioma cell implantation was opened again and two new cranial windows were opened near the previous cranial window, which used to label the ECM within the glioma at different area. Each cranial window was injected with 0.4 µL Atto 488 NHS dye in 1x PBS (0.25 mM) at a depth of 2 mm. After finishing the injection, the mouse was left in the stereotaxic frame for 0.5 h and then was ready for acute brain slice preparation.

##### Fluorescently labeling brain ECM of stroke mice

A cranial window was opened, centered at the coordinates of A/P +1.0 mm, M/L +1.5 mm. 0.4 µL Atto 488 NHS ester dye in 1x PBS (0.25 mM) was administered into the brain tissue via the cranial window with 0.8 mm injection depth. Then, the mouse was left in the stereotaxic frame for 0.5 h and ready for acute brain slice preparation.

#### Acute brain slice preparation

The acute brain slice (coronal) with 300 μm thickness was prepared using a vibratome (*Leica VT1200S*). The slicing process was performed in ice-cold low-Na^+^ low-Ca^2+^ high-Mg^2+^ artificial cerebrospinal fluid (ACSF) solution (slicing ACSF), which contains 72 mM NaCl, 2.5 mM KCl, 26 mM NaHCO_3_, 1.25 mM NaH_2_PO_4_, 0.5 mM CaCl_2_, 2 mM MgCl_2_, 24 mM glucose, and 75 mM sucrose (∼300 mOsm/L). Then, brain slices were transferred to a brain slice keeper (Automate Scientific, BSK2) with the recovery ACSF solution, which included 124 mM NaCl, 5 mM KCl, 26 mM NaHCO_3_, 1.25 mM NaH_2_PO4, 1.5 mM CaCl_2_, 1.3 mM MgCl_2_, and 10 mM glucose (∼300 mOsm/L), for 0.5 h at 34 °C and subsequently for 1 h at room temperature to recover brain slices’ activities. All ACSF buffers were bubbled with carbogen gas (95% O_2_ and 5% CO_2_).

#### Live brain ECM imaging

To maintain the acute brain slice’s activities during imaging, a perfusion system was built by coupling an open bath recording chamber (*Warner Instrument*, RC-26), a heating platform (PH-1, *Warner Instrument*), a peristaltic pump (*Sigma-Aldrich*, Z678406), and a vacuum system (*Warner Instrument*, 64-1940). The recording chamber and a #1.5 coverslip (*Fisher Scientific*) were put in the heating platform, which was incorporated into a microscope stage adapter (*Warner Instrument*, SA-20UU-AL). The heating platform was connected to a dual-channel temperature controller (*Warner Instrument*, TC-344C). The oxygenated recovery ACSF solution was flowed via the peristaltic pump and was further heated by a single inline solution heater (*Warner Instrument*, SH-27B). The perfusion rate was set as 2 mL/min, and the temperature was heated to 34 °C. Then, the acute brain slice was transferred to the recording chamber and was kept close to the coverslip using a slice anchor (*Warner Instrument*, SHD-26H/15). The brain ECM imaging was performed on an Olympus SD-OSR spinning disk confocal microscope, coupled with four laser lines, 405 nm, 488 nm, 561 nm, and 640 nm, and a CSU-W1 Yokogawa. The 100× silicone oil objective (NA 1.35) was used, achieving up to ∼50 microns imaging depth away from the coverslip surface. During the imaging acquisition, the ACSF perfusion was briefly paused for image acquisition to reduce motion artifacts. The brain ECM was visualized with the 488 nm laser excitation, and emission was collected by a bandpass filter 525/50 nm (FITC/Cy2 filter). The laser power and exposure time was adjusted to maximize the signal-to-noise ratio on the generated image. Each obtained image has a size of 1024 × 1024 pixels (pixel size 0.13 µm).

#### Live brain ECS imaging

Acute brain slices were prepared using the previous method and maintained in a perfusion system. All fluorophores for ECS imaging were listed in **Supplementary Table 1**. Before imaging, dyes were prepared in ACSF buffer with a concentration of 0.1 mM, followed by continual oxygen bubbling. Then, the fluorophores-ACSF buffer flowed into the recording chamber so that the brain slice could soak in the fluorophore solution, and the ECS of the brain slice would be filled with the fluorophore. The ECS of the acute brain slices was also imaged using the spinning disk microscope.

#### Fixed brain ECM imaging

The fixed brain ECM imaging was used for imaging dyes in the brain tissues in a large field-of-view (**Fig. 1c-d**, and **Supplementary Fig. 1**). Briefly, an acute brain slice (coronal) with 1 mm thickness that covered dyes injection area was prepared using the vibratome, followed by perfusion with oxygenated recovery ACSF solution (5 mL/min) for 30 minutes. Then, the brain slice was fixed with 4 % paraformaldehyde (PFA, *Thermo Fisher Scientific*, catalog NO. J19943.K2) in 1x PBS overnight at 4 °C and subsequently was quickly frozen with dry ice and transferred to the Cryostat (*Leica*, CM1850) for further slicing. 30 µm thickness coronal brain sections were collected on a glass slide and were washed with 1x PBS three times to remove the cryoprotectant agent. After that, the remaining water on the glass slide was removed with lab wipes. The glass slide and a #1.5 coverslip were glued together with a mounting medium (Fluoromount-G, *SouthernBiotech*., catalog NO.0100-01) and dried at room temperature overnight. Large field-of-view images were obtained using the 4x objective of the Olympus SD-OSR spinning disk, and entire brain slice images were obtained with a VS120 virtual slide microscope (Olympus).

#### Immunostaining

After labeling the brain ECM with Atto 488 NHS, a 1 mm acute brain slice was prepared on the vibratome. The brain slice was transferred to the perfusion chamber for 30 min ACSF perfusion with a rate of 5 mL/min. Then, the brain slice was fixed with 4 % PFA in 1x PBS overnight at 4 °C and subsequently immersed with 10 % sucrose (*Thermo Fisher Scientific*, catalog NO. 036508.30) solution (w/v, in 1x PBS) for 1 day, and 30 % sucrose solution (w/v, in 1x PBS) for another 1 day. The brain slice was quickly frozen with dry ice and was transferred to the Cryostat for further slicing. 30 µm thickness coronal brain sections were collected and were washed with 1x PBS three times. Brain sections were blocked with 1x PBS with 10% normal donkey serum (NDS, *SouthernBiotech*., catalog NO. 0030-01) and 0.2% Triton X-100 (TX, *Sigma-Aldrich*, product No. X100) for 2 h at room temperature. Afterward, brain sections were incubated with the primary antibody in 1x PBS with 2% NDS and 0.1% TX at 4 °C. After overnight incubation, brain sections were washed with 1x PBS three times and incubated with appropriate secondary antibodies in 1x PBS with 2% NDS and 0.1% TX for 2 h at room temperature. Both secondary antibodies were conjugated with Alexa Fluor 647 or Cy5. Then, brain sections were washed with 1x PBS three times, mounted on a #1.5 coverslip with mounting medium, and let dry overnight. All details of fluorescent marker, and primary or secondary antibodies were listed in **Supplementary Table 2**.

In order to stain neurons in brain slices, NeuroTrace™ 435/455 (Nissl staining, *ThermoFisher Scientific*, catalog NO. N21479) was utilized. Typically, brain slices were incubated with NeuroTrace™ 435/455 in 1x PBS (1:200 dilution) for 20 minutes following antibody staining. The brain slice was then washed twice with 1x PBS for 20 min and 2 h, respectively.

Low and high-resolution brain section images were obtained from the SD-OSR spinning disk microscope using the 4x (pixel size ∼3.30 µm) and 100x objective. The brain ECM was excited with a 488 nm laser, and antibodies staining was excited with a 640 nm laser. Emissions were filtered with two different bandpass filters in the SD-OSR microscope. **Fig. 4b-c** was obtained from the VS120 virtual slide microscope (Olympus), utilizing a SOLA-SE-II solid-state white light as the excitation source. Emissions were filtered with three bandpass filters, 49002 - ET - EGFP (FITC/Cy2), 49004 - ET - Cy3/TRITC, and 49006 - ET - Cy5.

#### Images analysis

Images processing was performed using *Fiji* (NIH). The look-up tables (LUT) were green for images collected from FITC/Cy2 channel, magenta for images collected from CY3/TRITC channel, and red for images collected from the Cy5 channel. Mean area fluorescence intensity in a region of interest (ROI) was quantified using the ‘Measure’ function. Fluorescence intensity profiles were generated by drawing a 3-pixel width line in the image and applying the ‘Plot Profile’ function. The colocalization between two images was evaluated using the *Imaris* software, and no image threshold was applied. Colocalization was quantified using the Pearson correlation coefficient (PCC, -1 to 1).

#### Estimation of dye diffusion volume

Atto 488 NHS, Cy3b NHS, and Atto 655 NHS were injected into the cortex as described in “Fluorescently labeling of brain ECM of wild-type C57BL/6 mice”, and fixed brain slices (30 µm thickness) covering the entire dye-labeled area were prepared. Each slice was viewed through a low magnification objective (4x) on the spinning disk confocal microscope. The intensity threshold for each raw confocal image was set above the background intensity, allowing the image to be binarized using Fiji’s ‘Make binary’ function. The binarized area for each image was determined using ‘Measure’ function, and the diffusion area of dye in each slice was calculated by multiplying the binarized area by the thickness of the slice (30 µm). The 3D diffusion volume was obtained by summing the diffusion areas for each slice.

#### Estimation of the ECM volume fraction

The ECM volume fraction (*α*) is defined as *α* = *V*_ECM_/*V*_Total_, where *V*_ECM_ represents the volume of ECM and *V*_Total_ is the total brain volume. To extract the ECM structure from the raw image, we employed the interactive wavelet-based segmentation Fiji plugin, ‘SpineJ’,^38^ adopting the second and third wavelet coefficients with a factor of 3. A white-black image was generated, which was then binarized using Fiji’s ‘Make Binary’ function. The binarized image was subjected to the ‘Measure’ function, where the ‘area%’ parameter represented the ECM volume fraction (*α*). Specifically, *α* was calculated as the number of pixels for the ECM structure in the binarized image divided by the total number of pixels in the binarized image.

#### Statistical analysis

The normality of data was determined first. Non-normal data were shown as medians with interquartile ranges, while normally distributed data were shown as mean with standard deviation (mean ± SD). Statistical significance analysis was carried out using appropriate parametric or non-parametric tests with a predetermined significance level of 5%. All statistical data analyses were done in Graphpad Prism. The sample size (n) and P values were indicated in the figure legend. Asterisks in figures indicate p values as follows: * p < 0.05, ** p < 0.01, *** p < 0.001.

## Acknowledgments

This study was partially sponsored by National Science Foundation (NSF) under grant number 2123971, National Institutes of Health (NIH) under grant number RF1NS110499, and McDermott Professorship at the University of Texas at Dallas.

## Author contributions

X.G. and Z.Q. generated the idea, and X.G. performed the experiment and analyzed the data. X.X. performed the IHC staining. X.X., Q.C., H.X., X.C., and X.-F.G., assisted in experiments and analyzed the data. Y.H., Y.Y., and R.B. help analyzed results. Z.Q. supervised the project, analyzed the data, and discussed the results. X.G., X.X., and Z.Q., wrote the paper. All authors revised the manuscript and have approved the final version.

## Conflict of Interest

The authors declare no conflict of interest.

## Data availability

All data associated with this study are present in the paper or the Supplementary Materials.

## Notes

### Competing Interest Statement

The authors have declared no competing interest.

