## Supplemental for "pan-ECM: live brain extracellular matrix imaging with protein-reactive dye"

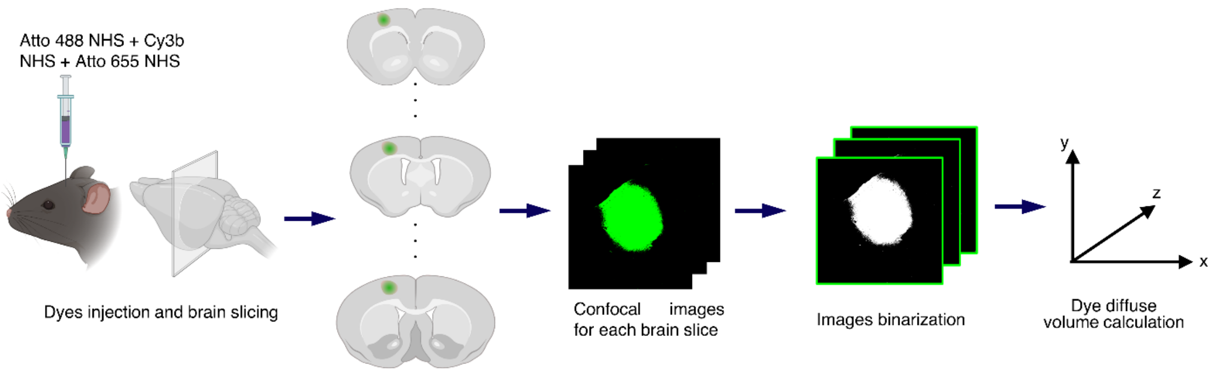


**Fig. S1 The calculation of dyes diffusion volume within brain slices.** Following labeling of the brain slice with NHS ester functionalized dye, fixed brain slices (30 µm in thickness) covering the dye-labeled area were prepared. To acquire raw confocal images, each slice was viewed through a low magnification objective (4x) on a confocal microscope. Afterward, raw confocal images were binarized, with the white area in the binary image being considered the dye diffusion area. For the 3D diffusion volume, it is the sum of the diffusion areas for each slice.

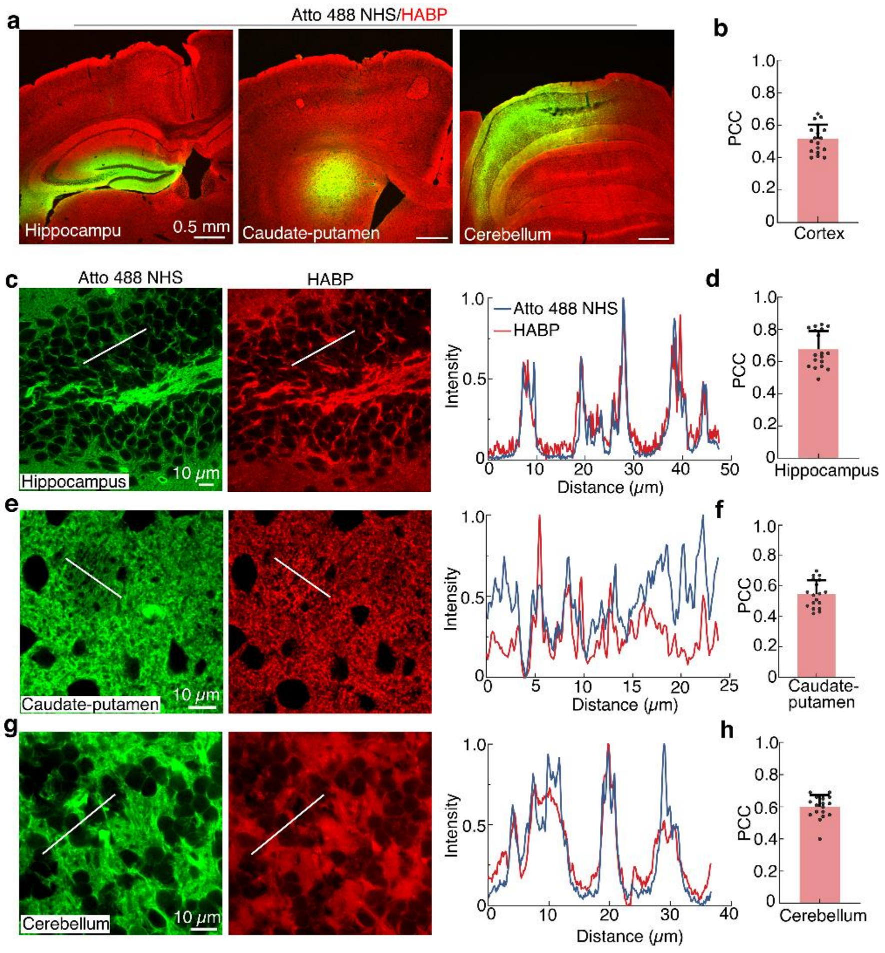


**Fig. S2 Colocalization analysis between Atto 488 NHS labeling and HABP staining**. (**a**) Large field-of-view images with Atto 488 NHS labeling and HABP staining. (**c, e,** and **g**) Confocal images of the extracellular structure in the hippocampus (**c**), caudate-putamen (**e**), and cerebellum (**g**) with Atto 488 NHS labeling and HABP staining; right: fluorescence intensity profiles along the white lines in panel c, e, and g, showing good colocalization between Atto 488 NHS labeling and HA staining. (**b**, **d**, **f**, and **h**) Pearson's coefficient values for evaluating the colocalization between Atto 488 NHS labeling and HABP staining in different brain regions.

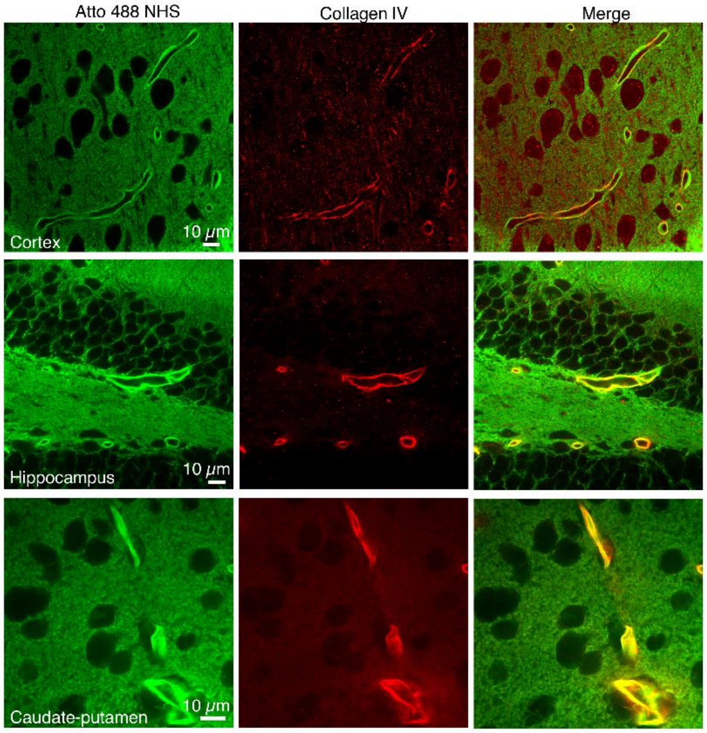


**Fig. S3 pan-ECM shows good labeling of the BM in the cortex, hippocampus, and caudate-putamen.** Left: Confocal images of extracellular structure with Atto 488 NHS labeling; middle: collagen IV staining; right: merging left and middle images.

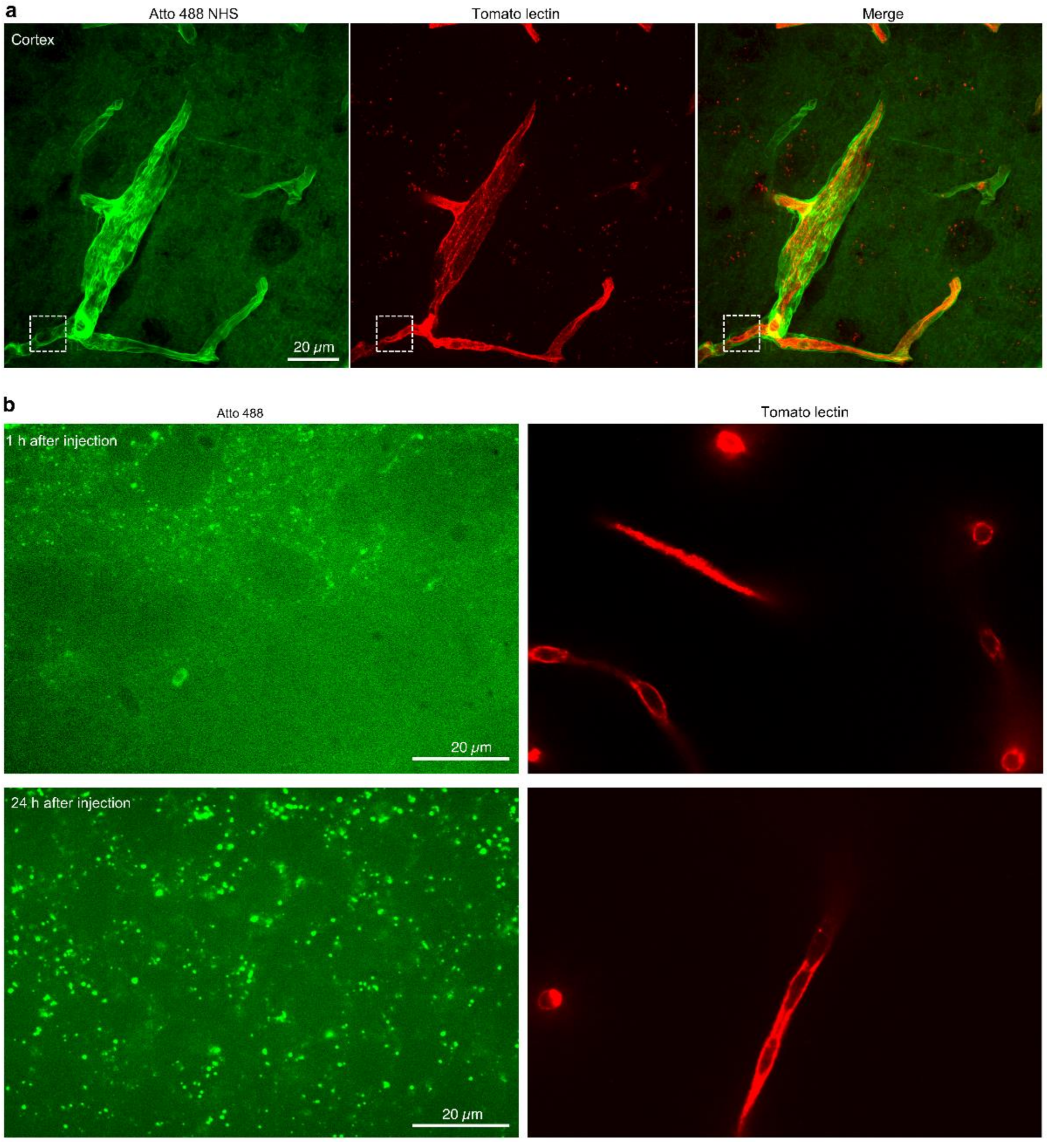


**Fig. S4 pan-ECM only labels the outside of the vascular lumen via covalent conjugation. (a)** Left, the confocal image of the extracellular structure in the brain cortex with Atto 488 NHS labeling; Middle, the vascular lumen was labeled with tomato lectin-Dylight 649; Right: the merge of left and middle images. Images from dashed boxes were shown in Fig. 2e. (**b**) Left: the confocal images of the extracellular structure in the hippocampus labeled with Atto 488 (no protein conjugation dye), following intracerebral injection at 1-hour and 24-hour intervals; right: the vascular lumen was labeled with tomato lectin-Dylight 649. These findings suggest that nonspecific binding precludes the deposition of Atto 488 dye in the basement membrane.

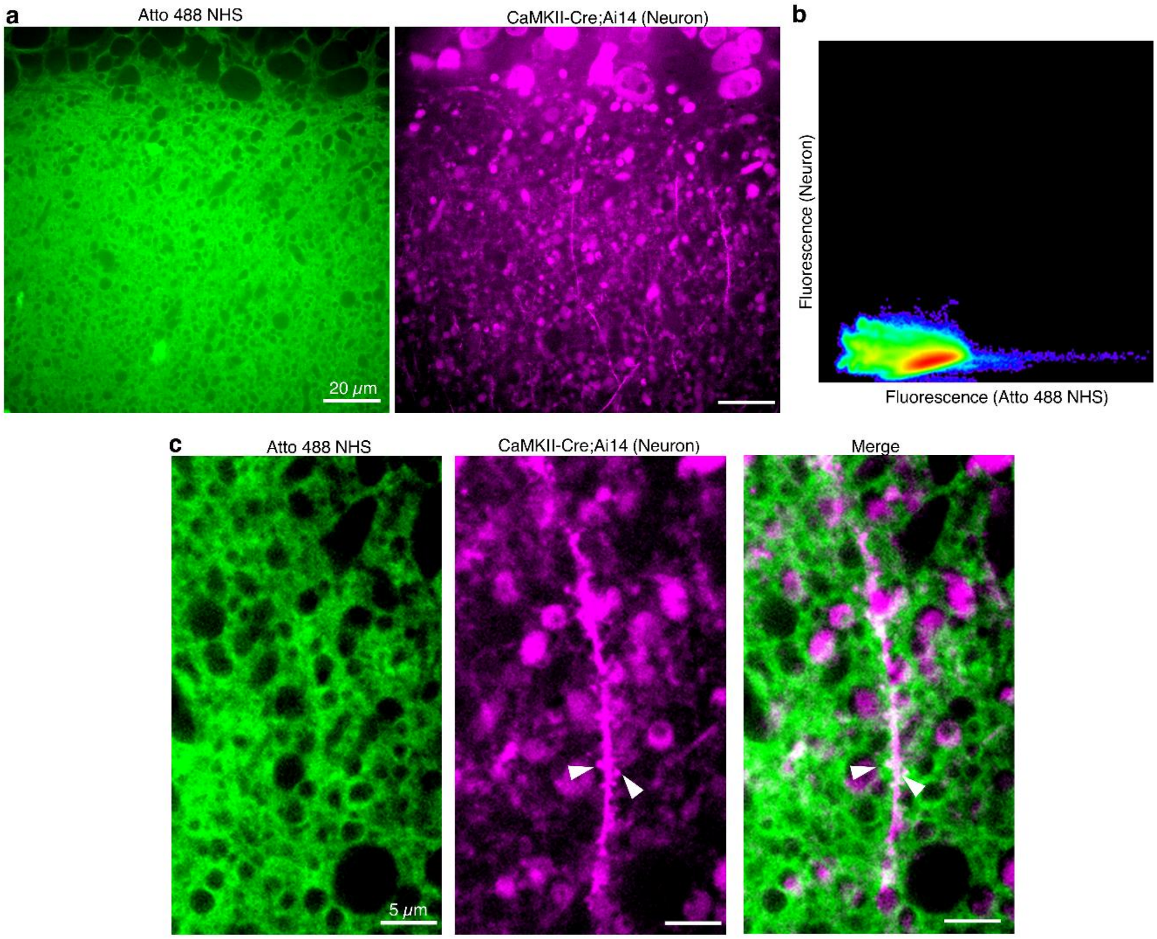


**Fig. S5 pan-ECM labels peri-neuronal matrix without neuronal uptake.** (**a**) Confocal images of (left) the extracellular structures with Atto 488 NHS labeling, (right) hippocampus neuron of the CaMKII -Cre:Ai14 mouse. (**b**) Scatter plot showing the rare colocalization of the Atto 488 NHS and neuron channels in panel a (Pearson's coefficient = 0.14). (**c**) The closed-up images of Atto 488 NHS labeled structures and neuron axons. Two dendritic spines were marked with white arrows.

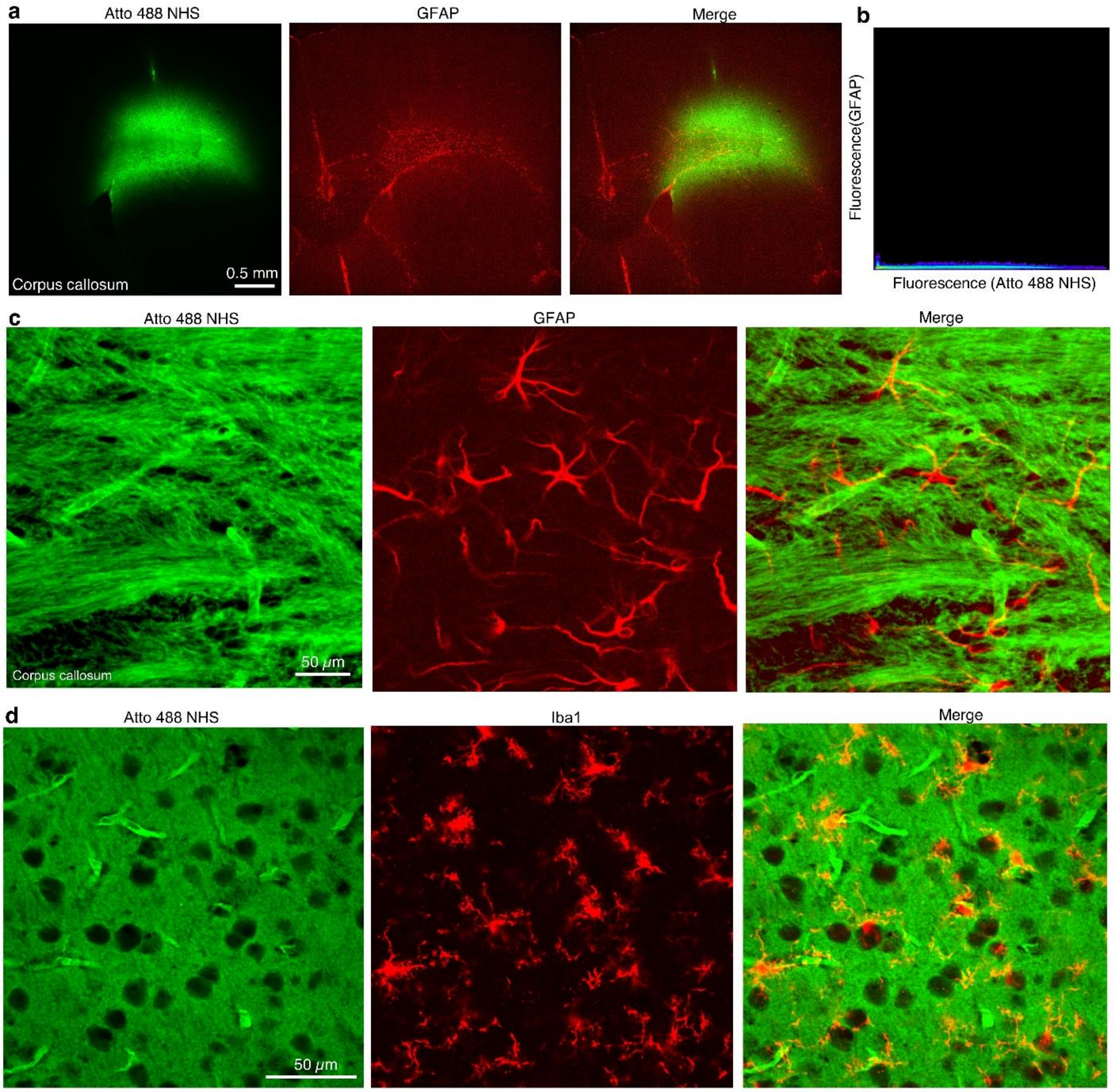


**Fig. S6 pan-ECM labeling does not lead to astrocyte and microglia uptake** (**a**) Confocal imaging on fixed brain slice with Atto 488 NHS labeling of extracellular structures and GFAP staining. (**b**) Scatter plot showing the rare colocalization of the Atto 488 NHS and GFAP channels (Pearson's coefficient = 0.08), showing no astrocytes uptake of Atto 488NHS. (**c**) The closed-up images of Atto 488 NHS labeled extracellular structures and GFAP staining. (**d**) Confocal images of the Atto 488 NHS labeled extracellular structures in the cortex and Iba1 staining (red), showing no microglia uptake of Atto 488NHS.

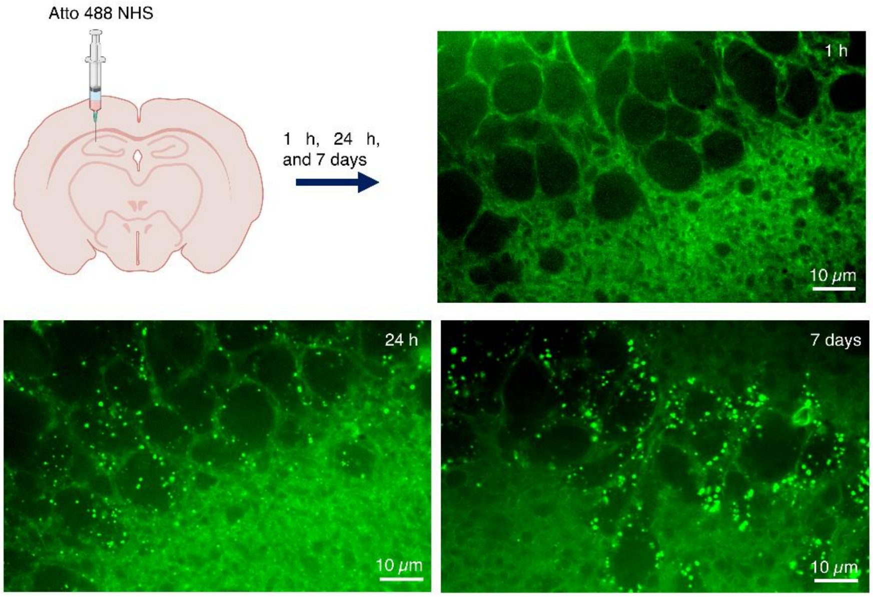


**Fig. S7 pan-ECM allows acute ECM labeling and can be taken up by hippocampal neurons at 24 hours.** Left: Schematic illustration of administrating Atto 488 NHS to the hippocampus to trace the endocytosis of Atto 488 dye; right: confocal images of Atto 488 NHS in the hippocampal neuron at 1 h, 24 h, and 7 days.

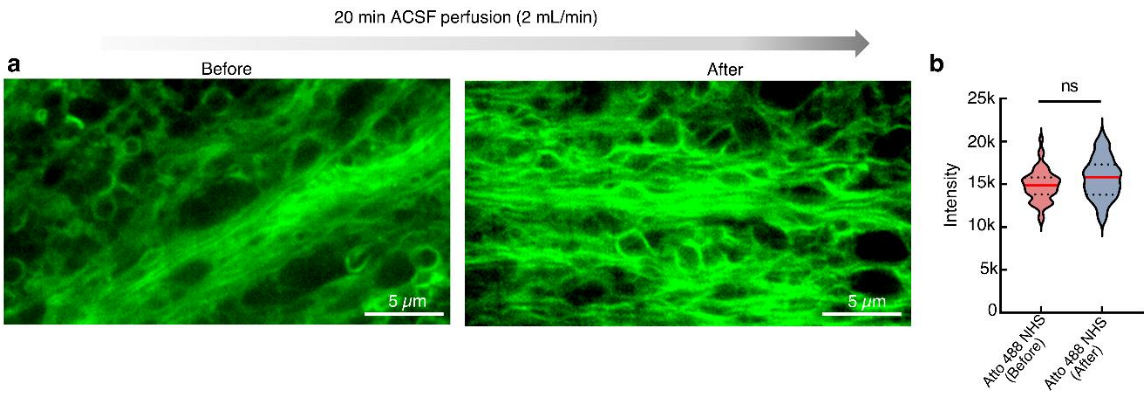


**Fig. S8 pan-ECM using NHS functionalized dye shows wash-resistant labeling**. (**a**) Live imaging of the ECM (labeled with Atto 488 NHS) before and after a 20 min ACSF buffer perfusion (2 mL/min). (**b**) Fluorescence intensity distributions of confocal images show the wash resistance for pan-ECM with Atto 488 NHS labeling. The errors indicate mean ± SD (n = 50 images from 6 brain slices, 3 mice); ns: P > 0.05, unpaired t-test.

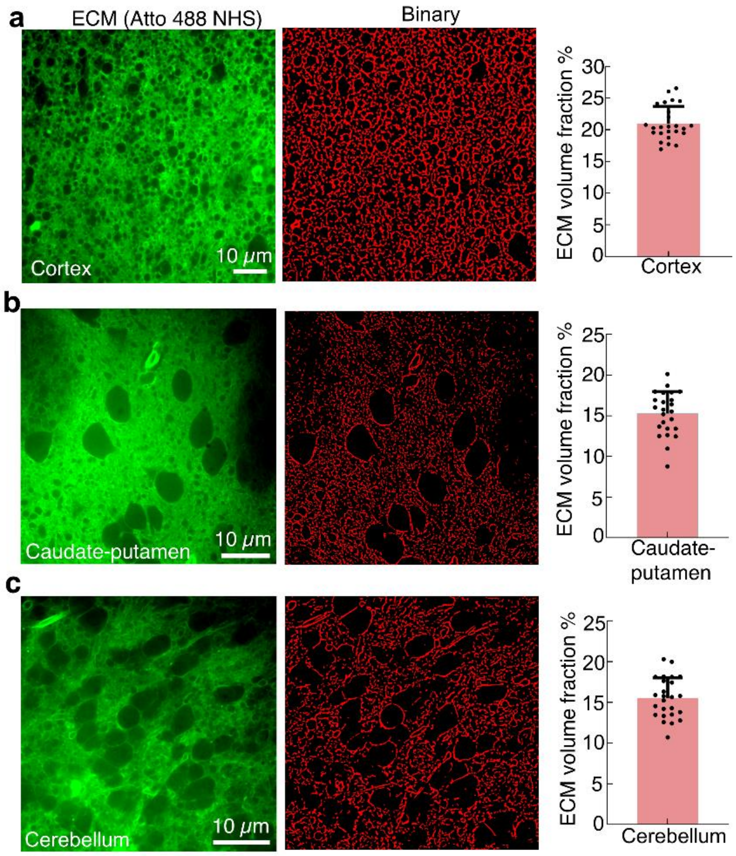


**Fig. S9 ECM volume fraction analysis across different brain regions.** Left: the confocal image of the brain ECM of (**a**) cortex, (**b**) caudate-putamen, and (**c**) the cerebellum in live tissue by pan-ECM labeling; middle, the binary of the confocal image; right, the ECM volume fraction distribution, mean ± SD. n = 25 images, for the cortex, n = 24 images for the caudate-putamen, and n = 25 mages for the cerebellum. Both images were obtained from 3 mice.

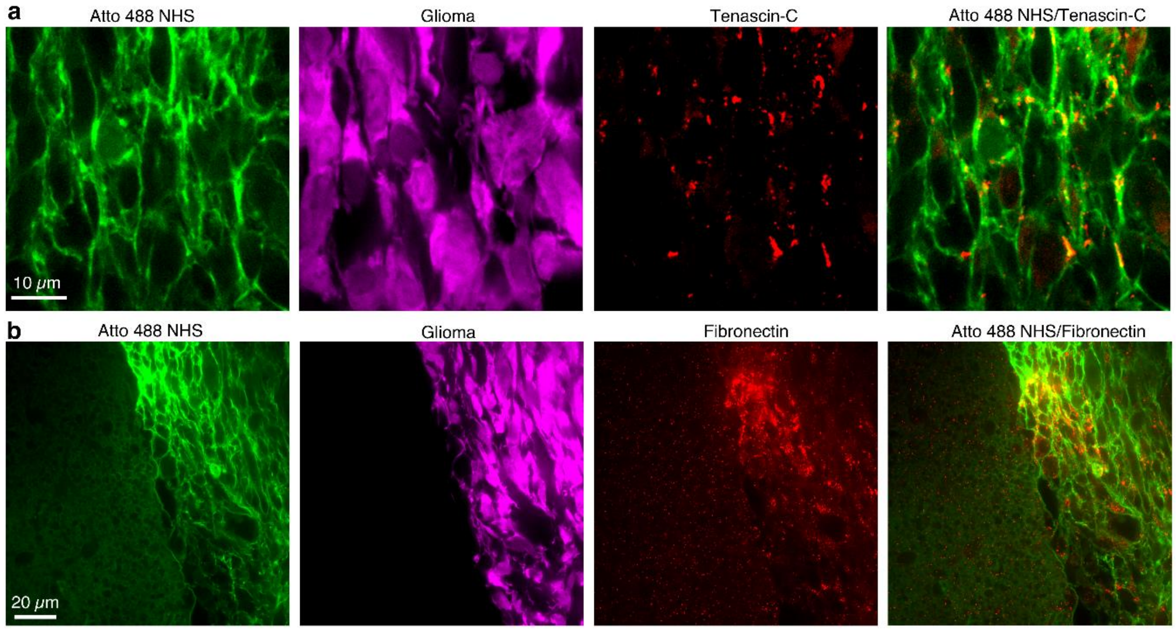


**Fig. S10 pan-ECM labels tenascin C and fibronectin in brain glioma**. Confocal images of the extracellular structure in the glioma with Atto 488 NHS labeling (green), glioma cell (magenta), antibodies staining (red), and the merge of images from green and red channels. (**a**) Tenascin-C, (**b**) fibronectin.

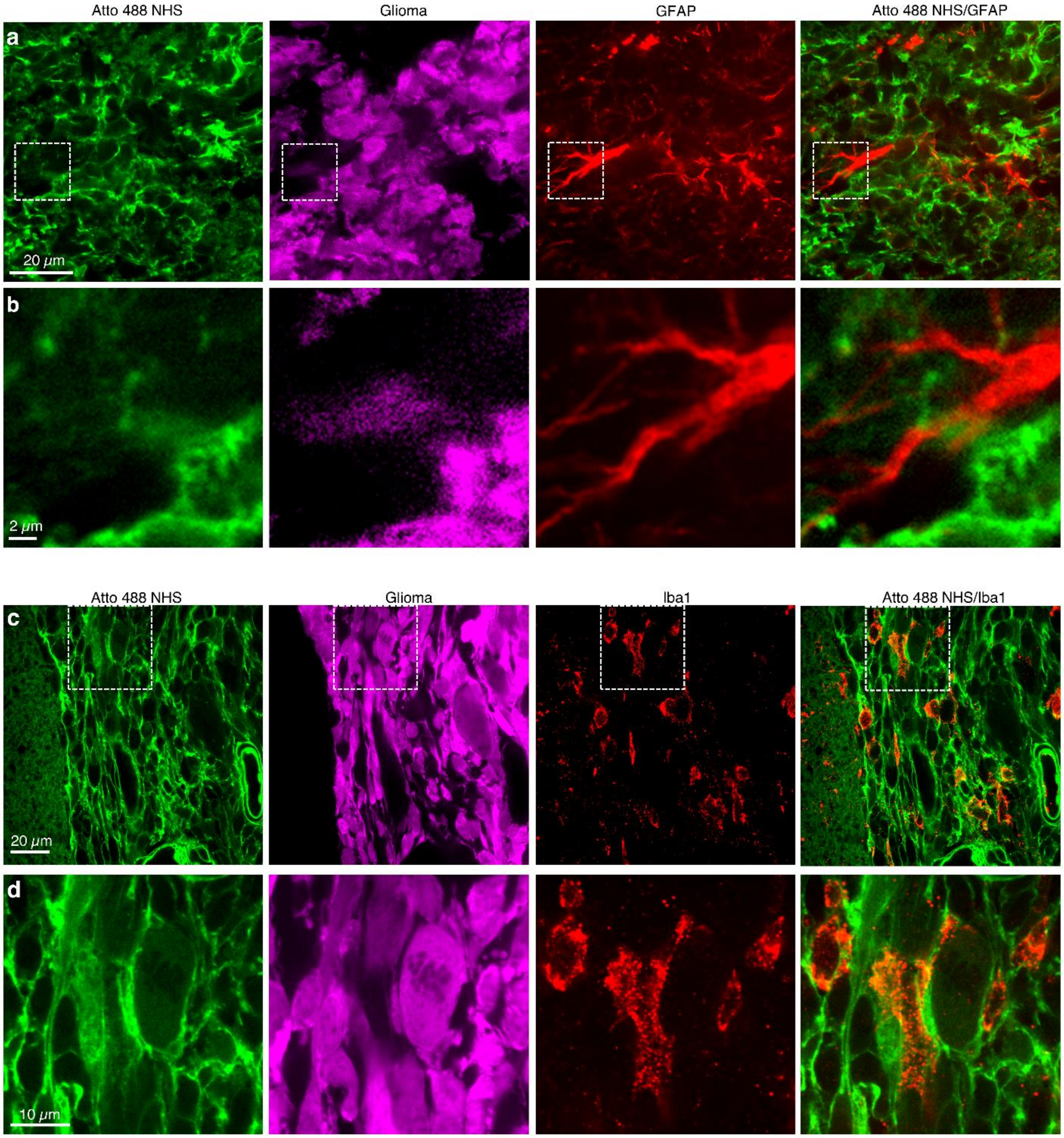


**Fig. S11 pan-ECM labeling of glioma ECM shows no cellular uptake of Atto 488 NHS dye within glial cells.** Confocal images of the extracellular structure in the glioma with Atto 488 NHS labeling (green), glioma cell (magenta), and antibodies staining (red). (**a** and **b**) GFAP staining for astrocytes, (**c** and **d**) Iba1 staining for microglia. Panels b and d display magnified images of the dashed boxes shown in panels a and c, respectively.

### Table S1 The summary of fluorescent dyes for brain extracellular matrix imaging

| Fluorophore | Vendor | Molecular structure | hydrophobic /hydrophilic | QY | Aggregation |
|---|---|---|---|---|---|
| Atto 488 | *Sigma-Aldrich* (41051) | 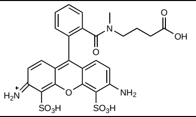 | Hydrophilic | 0.8 | No |
| Calcein | *Alfa Aesar* (L10255) | 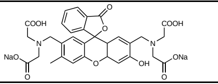 | Hydrophilic | 0.78 | No |
| Cy3b | *AAT Bioquest* (940) | 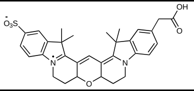 | Moderately hydrophilic | 0.58 | No |
| Atto 565 | *Sigma-Aldrich* (75784) | 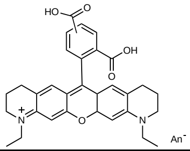 | Moderately hydrophilic | 0.9 | Aggregation |
| Atto 655 | *Sigma-Aldrich* (93711) | 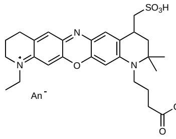 | Hydrophilic | 0.3 | No |
| Cy5 | *AAT Bioquest* (150) | 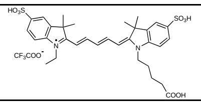 | Hydrophobic | 0.27 | Aggregation |
| Atto 647N | *Sigma-Aldrich* (04507) | 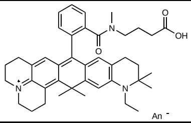 | Hydrophobic | 0.65 | Aggregation |
| Alexa Fluor 647 | *ThermoFisher Scientific*, (A33084) | 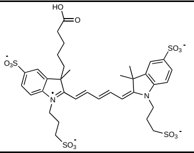 | Hydrophilic | 0.33 | Aggregation |

Atto dyes' properties were adopted from ATTO-TEC GmbH or Sigma-Aldrich. Cy3b and Cy5 dye properties were obtained from the AAT Bioquest. Calcein dye properties were obtained from Alfa Aesar. Alexa Fluor 647 properties were obtained from ThermoFisher Scientific.

**Table S2 The summary of primary antibodies and secondary antibodies for immunostaining**

| Targets | Primary antibody | Vendor | Secondary antibody | Vendor |
|---|---|---|---|---|
| Hyaluronan binding protein | Biotinylated hyaluronan binding protein | *Amsbio LLC* (AMS.HKDBC41) | Cy5-Streptavidin | *Invitrogen* (SA1011) |
| Perineuronal nets | Biotinylated wisteria floribunda lectin | *Vector* (B-1355-2) | Cy5-Streptavidin | *Invitrogen* (SA1011) |
| Chondroitin sulfate proteoglycans | Monoclonal anti-chondroitin sulfate antibody | *Sigma-Aldrich* (C8035) | Donkey anti-mouse IgG antibody, Alexa Fluor™ 647 | *ThermoFisher* (A-31571) |
| Collagen IV | Collagen IV polyclonal Antibody | *Invitrogen* (PA1-85320) | Donkey anti-Rabbit IgG antibody, Alexa Fluor™ 647 | *ThermoFisher* (A-31573) |
| Glial fibrillary acidic protein (GFAP, astrocyte) | GFAP rabbit polyclonal antibody | *ThermoFisher* (RB-087-A0) | Donkey anti-Rabbit IgG antibody, Alexa Fluor™ 647 | *ThermoFisher* (A-31573) |
| Ionized calcium-binding adapter molecule1(Iba1 , microglia) | Anti-Iba1 antibody | *Abcam* (ab5076) | Donkey anti-goat IgG antibody, Alexa Fluor™ 647 | *ThermoFisher* (A-21447) |
